# Gene Expression Elucidates Functional Impact of Polygenic Risk for Schizophrenia

**DOI:** 10.1101/052209

**Authors:** Menachem Fromer, Panos Roussos, Solveig K Sieberts, Jessica S Johnson, David H Kavanagh, Thanneer M Perumal, Douglas M Ruderfer, Edwin C Oh, Aaron Topol, Hardik R Shah, Lambertus L Klei, Robin Kramer, Dalila Pinto, Zeynep H Gümüş, A. Ercument Cicek, Kristen K Dang, Andrew Browne, Cong Lu, Li Xie, Ben Readhead, Eli A Stahl, Mahsa Parvisi, Tymor Hamamsy, John F Fullard, Ying-Chih Wang, Milind C Mahajan, Jonathan M.J. Derry, Joel Dudley, Scott E Hemby, Benjamin A Logsdon, Konrad Talbot, Towfique Raj, David A Bennett, Phil L De Jager, Jun Zhu, Bin Zhang, Patrick F Sullivan, Andrew Chess, Shaun M Purcell, Leslie A Shinobu, Lara M Mangravite, Hiroyoshi Toyoshiba, Raquel E Gur, Chang-Gyu Hahn, David A Lewis, Vahram Haroutonian, Mette A Peters, Barbara K Lipska, Joseph D Buxbaum, Eric E Schadt, Keisuke Hirai, Kathryn Roeder, Kristen J Brennand, Nicholas Katsanis, Enrico Dominici, Bernie Devlin, Pamela Sklar

## Abstract

Over 100 genetic loci harbor schizophrenia associated variants, yet how these common variants confer risk is uncertain. The CommonMind Consortium has sequenced dorsolateral prefrontal cortex RNA from schizophrenia cases (n=258) and control subjects (n=279), creating the largest publicly available resource to date of gene expression and its genetic regulation; ∼5 times larger than the latest release of GTEx. Using this resource, we find that ∼20% of the schizophrenia risk loci have common variants that could explain regulation of brain gene expression. In five loci, these variants modulate expression of a single gene: *FURIN, TSNARE1, CNTN4, CLCN3 or SNAP91*. Experimentally altered expression of three of them, *FURIN*, *TSNARE1*, and *CNTN4*, perturbs the proliferation and apoptotic index of neural progenitors and leads to neuroanatomical deficits in zebrafish. Furthermore, shRNA mediated knock-down of *FURIN1* in neural progenitor cells derived from human induced pluripotent stem cells produces abnormal neural migration. Although 4.2% of genes (N = 693) display significant differential expression between cases and controls, 44% show some evidence for differential expression. All fold changes are ≤ 1.33, and an independent cohort yields similar differential expression for these 693 genes (r = 0.58). These findings are consistent with schizophrenia being highly polygenic, as has been reported in investigations of common and rare genetic variation. Co-expression analyses identify a gene module that shows enrichment for genetic associations and is thus relevant for schizophrenia. Taken together, these results pave the way for mechanistic interpretations of genetic liability for schizophrenia and other brain diseases.

---

The human brain is complicated and not well understood. Seemingly straightforward fundamental information such as which genes are expressed therein and what functions they perform are only partially characterized. To overcome these obstacles, we established the CommonMind Consortium (CMC; www.synpase.org/CMC), a public-private partnership to generate functional genomic data in brain samples obtained from autopsies of cases with and without severe psychiatric disorders. The CMC is the largest existing collection of collaborating brain banks and includes over 1,150 samples. A wide spectrum of data is being generated on these samples including regional gene expression, epigenomics (cell-type specific histone modifications and open chromatin), whole genome sequencing, and somatic mosaicism.

Schizophrenia (SCZ), affecting roughly 0.7% of adults, is a severe psychiatric disorder characterized by abnormalities in thought and cognition (*1*). Despite a century of evidence establishing its genetic basis, only recently have specific genetic risk factors been conclusively identified, including rare copy number variants (*2*) and >100 common variants (*3*). However, there is not a one-to-one Mendelian mapping between these SCZ risk alleles and diagnosis. Instead, SCZ is truly complex and appears to result from a myriad of genetic variants exerting small effects on disease risk (*4*, *5*), conforming closely to a classical polygenic model (*6*). The available data are incomplete but implicate synaptic components, including calcium channel subunits and post-synaptic elements (*5*, *7*-*9*). A consequence of polygenic inheritance is that the small effect sizes of individual variants complicate characterization of the biological processes they influence, both at the level of particular genes and pathways.

Post-mortem gene expression studies of SCZ cases suggest subtle abnormalities in multiple brain regions including the prefrontal and temporal cortices, hippocampus, and several specific cell types (*10*). More than 50 gene expression studies of SCZ cases and controls have been reported, often of overlapping samples and mostly of modest scale (prior RNA sequencing studies evaluated only 5-31 cases, Supplementary data file 1). Results are often inconsistent and there are few replicated findings. These studies are probably underpowered to detect subtle effects that might be expected to arise as a result of this complex disease and within tightly regulated brain tissue (*11*), among other limitations of existing microarray-based gene expression studies (*12*, *13*).

RNA sequencing can accurately detect transcription at the gene and isoform level (*14*-*20*). We sequenced a cohort of SCZ and control subjects that is an order of magnitude larger than prior RNA sequencing studies. By applying state-of-the-art analytic methods and including genome-wide characterization of common variants, we generated a rich resource of the genetics of gene expression in the brain. This resource can serve as a useful catalogue of regulatory variants underlying the molecular basis of SCZ and other brain disorders. We use this resource to identify: (a) specific effects on gene expression of genetic variants previously implicated in risk; (b) genes showing a significant difference in expression between SCZ cases and controls; and (c) coordinated expression of genes implicated in SCZ. Our results shed light on the subtle effects expected from the polygenic nature of SCZ risk and thus substantially refine our understanding of the neurobiology of SCZ.

## Samples and sequencing

We generated RNA sequence data from post-mortem human dorsolateral prefrontal cortex (DLPFC; Brodmann areas 9 and 46) from brain banks at the Icahn School of Medicine at Mount Sinai, the University of Pennsylvania, and the University of Pittsburgh. To control for batch effects, multiple randomization steps were introduced and DNA and RNA isolation and library preparation were performed at one site (Supplementary Fig. 1A). Samples were genotyped on the Illumina Infinium HumanOmniExpressExome array (958,178 SNPs) and imputed using standard techniques with the 1000 Genomes Project as reference data (*21*). These genotypes were then used to detect SNPs that have an effect on gene expression (eQTLs, expression quantitative trait loci), to estimate ancestry of the samples, and to ensure sample identity across DNA and RNA experiments. Ethnicity was similar between cases and controls (Caucasian 80.7%, African-American 14.7%, Hispanic 7.7%, East Asian 0.6%, Supplementary Figs. 1B, C). As expected (*3*), SCZ cases inherited an increased number of common variant alleles previously associated with SCZ risk (p = 1.6 x 10^−8^, Supplementary Fig. 1D).

RNA sequencing was performed after depleting ribosomal RNA (rRNA). Following quality control, there were 258 SCZ cases and 279 controls. Fifty-five cases with affective disorder were included to increase power to detect eQTLs. The median number of paired end reads per sample was 41.6 million, with low numbers of rRNA reads (Supplementary Fig. 2). Following data normalization, 16,423 genes (based on Ensembl models) were expressed at levels sufficient for analysis, of which 14,222 were protein coding. Validation using PCR showed high correlation (r > 0.5) with normalized expression from RNA-seq for the majority of genes assessed (Supplementary Fig. 3). Gene expression measurement can be influenced by a number of variables; some are well documented (e.g., RNA integrity (RIN) and post-mortem interval (PMI)), but others may be unknown. We investigated known covariates by standard model selection procedures to find a good statistical model (Supplementary Fig. 4). Covariates for RIN, library batch, institution (brain bank), diagnosis, age of death, genetic ancestry, PMI, and sex together explained a substantial fraction (0.42) of the average variance of gene expression, and were thus employed to adjust the data for all analyses.

## Generation of a brain eQTL resource

To identify eQTLs, gene expression data from European-ancestry subjects (n=467) were adjusted for known and hidden variables detected by surrogate variable analysis (SVA) conditional on diagnosis but excluding ancestry (Supplementary Fig. 2 and 4). Adjusted expression levels were then fit to imputed SNP genotypes, covarying for ancestry and diagnosis, using an additive linear model implemented in MatrixEQTL (*22*). The model identified 2,154,331 significant cis-eQTLs, (i.e., within 1 Mb of a gene) at a false discovery rate (FDR) **≤** 5%, for 13,137 (80%) of 16,423 genes. Many eQTLs for the same gene were highly correlated, due to linkage disequilibrium, and 32.8% of eQTL SNPs (“eSNPs”) predict expression of more than one gene. Cis-eSNPs were enriched within genic elements and non-coding RNAs, particularly within 100 kb of the transcription start and end sites (*23*), and depleted in intergenic regions (Fig. 1A, B). As defined by GTEx (*24*), an “eGene” is a gene with at least one significant eSNP after strict correction for multiple marker testing for that gene. There were 8,427 eGenes at FDR **<** 5%, or 18 eGenes discovered per sample, consistent with a prediction from GTEx. We examined the enrichment of max-eQTLs (defined as the most significant eSNP per gene, if any) in predicted enhancer sequences derived from the Roadmap Epigenomics Consortium and ENCODE across 98 human tissues and cell lines (*25*). Cis-eQTLs were enriched for enhancer sequences present in brain tissues (Kolmogorov-Smirnov (KS) test versus non-brain: *D* = 1, p = 4.5 x 10-^6^), and the strongest enrichment is observed in DLPFC enhancers (*Z* = 9.5) (Fig. 1C).

**Figure 1.**
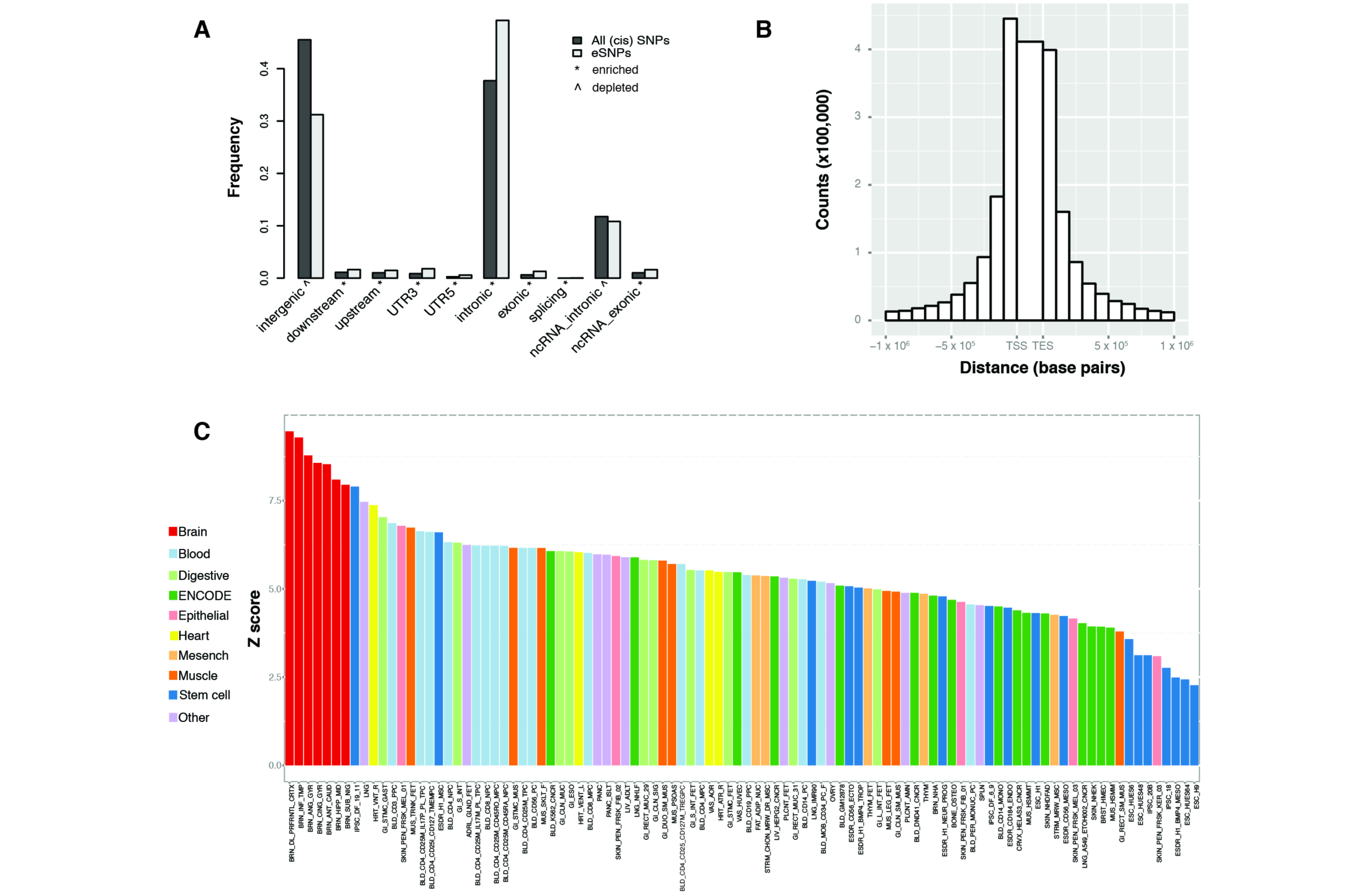
Enrichment of cis-eQTLs in regulatory and other genomic elements. **(A)** Enrichments of cis-eQTLs compared to all eQTLs in sequence-defined elements according to the Ensembl annotations implemented in the ANNOVAR (Annotate Variation, version 2014-07-14; Ensembl annotation) software (*71*). The y-axis illustrates the proportion of SNPs that belong to each category for significant cis-eQTLs (at FDR 5%) compared to all eSNPs that were considered for cis-interactions (within 1 Mb from expressed genes). The following categories are illustrated in the figure: exonic; intronic; upstream (variant overlaps 1 kb region upstream of transcription start site); downstream (variant overlaps 1 kb region downstream of transcription end site); UTR3 (variant overlaps a 3’ untranslated region); splicing (variant is within 2 bp of a splicing junction); ncRNA (variant overlaps a transcript without coding annotation in the gene definition followed by additional annotation for exonic, intronic, variants as described above); intergenic. (^) and (*) indicate significant depletion or enrichment for certain genic categories of cis-eQTLs compared to all eSNPs, respectively**. (B)** Enrichment of cis-eQTLs as a function of distance from the transcription start and end sites. **(C)** Enrichment of “max-cis-eQTLs” (single most associated eSNP per gene) within enhancer sequences across 98 human tissues and cell lines. Each bar represents the *Z* score for the overlap of max-cis-eQTLs with each enhancer compared to 1,000 sets of random SNPs matched with the max-cis-eQTLs, in terms of allele frequency, gene density, distance from the transcription start site, and density of tagSNPs due to linkage disequilibrium. Brain (red) shows significantly higher enrichment for eQTLs compared to non-brain tissues and cell lines (p = 4.5 x 10^−6^) and the strongest enrichment is observed in DLPFC enhancers.

To assess the utility of our much larger brain dataset, we compared previously reported DLPFC eQTLs to CMC-derived eQTL, estimating the proportion of non-null hypotheses *(TT_1_)* in CMC (*26*) and the number of additional eQTL found in CMC that were not detected in the other studies. GTEx v6 is the largest public dataset of eQTLs from DLPFC tissue (n = 92) assayed by RNA-seq; its replication in CMC is ***TT***_*1*_ = 0.98. Considering microarray-based eQTLs from the Harvard Brain Bank (*27*), BrainCloud (*28*), NIH (*29*), and the UK Brain Expression Consortium (UKBEC) (*30*), we estimated ***TT***_*1*_ to be 0.75, 0.70, 0.79, and 0.93, respectively, indicating that our results captured most eQTLs found in other independent samples. Replication was somewhat lower for a recent meta-analysis that included mixed several distinct brain regions (*31*) (**TT**_1_ = 0.62), and for eQTLs detected in blood (***TT***_*1*_ = 0.54) (*32*). We also derived eQTL for 279 DLPFC samples as part of the NIMH Human Brain Collection Core (HBCC) microarray data and found replication ***TT**_1_ = 0.77*. Moreover, concordance of the direction of allelic effect was high, with 93% of eQTL showing the same direction of effect when intersecting CMC eQTL (FDR **≤** 5%) with even a liberally defined set of HBCC eQTL (FDR **≤** 20%). In addition to containing the vast majority of eQTL found in the literature, the CMC sample finds a substantial number of genes with previously undetected eQTL (Table 1).

**Table 1:** Overlaps and differences between CMC and other publicly available eQTL resources. * Best eQTL per probeset reported ** Best eQTL per gene reported FDR **<** 5% used to define eQTL in all cohorts. eQTL for Brain Cloud, HBCC, HBTRC, NIH and UKBEC were computed as described in the supplement. eQTL for the Blood cohort, Brain MetaGanalysis and GTEx were downloaded from public resources. All eQTL resources represent prefrontal or frontal cortex except the Blood cohort (peripheral blood) and the Brain MetaGanalysis (metaGanalysis across multiple brain regions). The UNION set was derived by including all unique eQTL from all 8 cohorts.

| Cohort | Sample Size | Study PMID/GEO ID/dbGaP ID | Number of cis eQTL | Comparison cohort eQTL genes compared to CMC eQTL |  |  |  | Genes w/ eQTL in CMC but not in comparison cohort |
| --- | --- | --- | --- | --- | --- | --- | --- | --- |
| | | | | Proportion of non-null hypotheses ($\pi_1$) in CMC | Unique Genes with eQTL | eQTL Genes Expressed in CMC | Genes with eQTL in CMC | |
| Blood eQTL | 2494 twins | 24728292 | 9640* | 0.54 | 9533 | 8108 | 6794 | 5052 |
| Brain Cloud | 108 | GSE30272 | 374223 | 0.7 | 6199 | 5386 | 4666 | 7180 |
| Brain Meta-analysis | 424 | 25290266 | 3520** | 0.62 | 3503 | 2806 | 2507 | 9339 |
| GTEx PFC | 92 | 25954002 | 173026 | 0.98 | 1922 | 1326 | 1284 | 11853 |
| HBCC | 279 | phs000979.v1.p1 | 788338 | 0.77 | 7514 | 6785 | 5862 | 7275 |
| HBTRC | 146 | GSE44772 | 531400 | 0.75 | 6473 | 5186 | 4555 | 7291 |
| NIH | 145 | GSE15745 | 105735 | 0.79 | 2127 | 2057 | 1851 | 9995 |
| UKBEC | 134 | 25174004 | 52593 | 0.93 | 808 | 618 | 546 | 11300 |
| **UNION** |  |  | **1573706** | **0.7** | **16568** | **12644** | **10544** | **2593** |

\* Best eQTL per probeset reported
\*\* Best eQTL per gene reported

The patterns of results should be different for “trans-eQTLs”, i.e., SNPs correlated with expression of a gene beyond 1 Mb of its genomic location. Trans-eQTLs incur a greater penalty for multiple testing, require greater power for detection, and are thus more susceptible to false positives and less likely to replicate than cis-eQTL. Nevertheless, the data supported 45,453 significant trans-eQTL at FDR ≤ 5%, of which 20,288 were also cis-eQTL SNPs for local genes, and 34% predicted expression of more than one distant gene. The proportion of trans eQTL in CMC that replicate in HBCC is 18.6% (both FDR ≤ 5%). The proportion of HBCC trans eQTL that replicate in CMC is 29.7%. Enrichment of trans-eQTLs with brain enhancers was not observed (data not shown), though enrichment in genic regions and depletion in intergenic regions was observed, particularly when restricting to trans eQTL **≥** 10 Mb from the gene location. We used similar techniques to derive isoform expression quantitative trait loci (isoQTLs). Those results are described in Supplementary Information.

## eQTL signatures at SCZ risk loci point to specific genes

A hallmark of polygenic inheritance is that individual SNPs confer small effects on risk. For some risk SNPs, perhaps the majority, their impact could be mediated through effects on gene expression. Indeed, GWAS SNPs associated with SCZ risk occur more often than expected by chance in cis-regulatory functional genomic elements, such as enhancers or eQTL SNPs (*7*, *33*-*35*). Yet, GWAS loci typically contain many genes, and SNPs therein are often highly correlated via linkage disequilibrium, so that assigning a biological role for a particular risk SNP has been difficult. Here, we leverage CMC-derived eQTL to relate SCZ risk variants to expression of specific genes.

Of the 108 SCZ GWAS loci previously reported (*7*), 73 harbor cis-eQTL SNPs for one or more genes (FDR **≤** 5%). However, the simple presence of an eQTL does not imply disease causality (*36*). We used Sherlock (*37*), a Bayesian approach that prioritizes consistency between disease association and eQTL signatures in GWAS loci, to identify genes likely to contribute to SCZ etiology. While Sherlock evaluated genes across the genome, we only evaluated genes within the 108 SCZ GWAS loci because SNPs in these loci showed genome-wide significant association with SCZ; thus, in essence, we fine mapped these loci. The results suggested that GWAS risk and eQTL association signals co-localized for 84 genes in 30 of these loci (adjusted p < 0.05; Supplementary Fig. 5A, data file 2). After removing genes where additional evaluation indicated lack of consistency (Supplementary Fig. 6B), there were 33 genes highlighted in 18 of the 108 GWAS loci (data file 2). Genes found to have variants affecting risk for autism are often found enriched for variation affecting risk for SCZ; indeed, compared to other genes with eQTL in the GWAS loci, these 33 genes are more enriched for nonsynonymous de novo mutations in autism (fold enrichment = 2.4, p_corrected_ = 0.03), although not for SCZ, intellectual disability, or epilepsy.

Repeating the analyses using isoform-level eQTLs (isoQTL) identified nine genes in eight GWAS loci, with all but three genes already identified in the gene-level analysis (data file 2). Combining the gene and isoform data, 20 of 108 GWAS loci (19%) had evidence suggesting that mis-regulated gene expression could, in part, explain the genetic association with schizophrenia: 18 cis-QTL loci (cis-eQTL for 33 genes + 2 genes with cis-isoQTL), one locus implicated only by cis-isoQTL (*SNX19*), and one trans-eQTL association for *IMMP1L* at a GWAS locus on chr7. We discuss other genes identified by Sherlock in the Supplement.

Of the 19 GWAS loci harboring SCZ-associated cis-eQTLs, eight involved only a single gene (i.e., no additional gene with relaxed adjusted Sherlock p < 0.5): furin (*FURIN*, down-regulated by risk allele), t-SNARE domain containing 1 (*TSNARE1*, up), contactin 4 (*CNTN4*, up), voltage-sensitive chloride channel 3 (*CLCN3*, up), synaptosomal-associated protein of 91 kDa (*SNAP91*, up), ENSG00000259946 (up), ENSG00000253553 (down), and the ENST00000528555 isoform of sorting nexin 19 (*SNX19*, down) (Fig. 2 and Supplementary Fig. 5B and 6A). For functional follow-up, we focused on the five single-gene loci encoding known proteins implicated at the gene level. First, we replicated these eQTL in the Religious Orders Study and Memory and Aging Project (ROS/MAP) (*38*), with unpublished human DLPFC RNA sequencing data (n=461). The most significant GWAS SNP was also a significant eQTL with the same direction of effect as in CMC for *FURIN* (rs4702: p = 1 x 10^−6^), *CLCN3* (rs10520163: p = 9 x 10^−6^), and *SNAP91* (rs3798869: p = 3 x 10^−4^); *TSNARE1* (rs4129585: p = 0.057) and *CNTN4* (rs17194490: p = 0.07) also had alleles in the same direction of effect as in CMC but did not reach significance.

**Figure 2.**
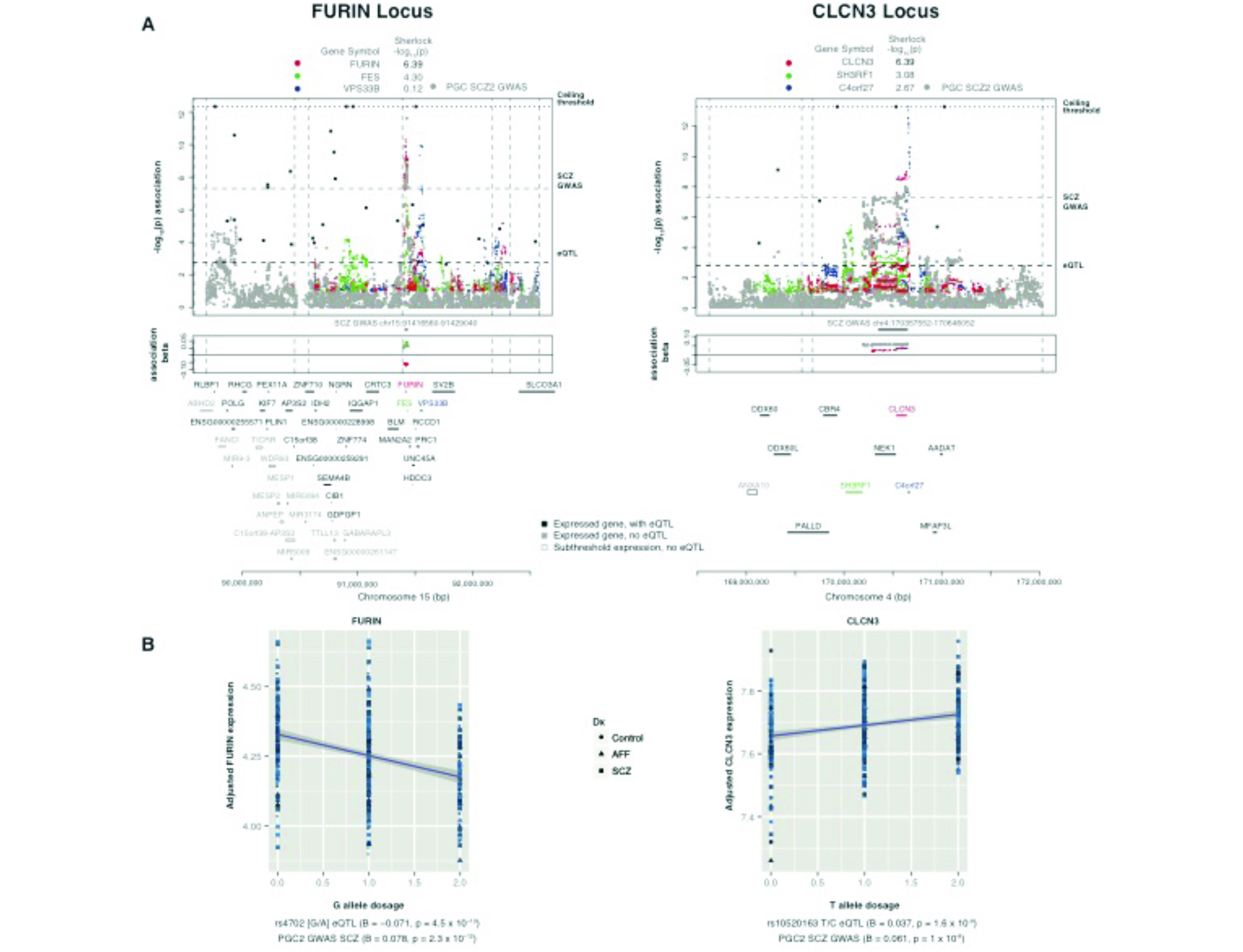
Overlap of GWAS for schizophrenia with eQTL in the DLPFC. **(A)** eQTL association profiles across two representative SCZ GWAS loci on chromosomes 15 and 4, respectively. SNP-level associations are plotted for the SCZ GWAS (gray), and cis-eQTL association profiles for genes with Sherlock p_corrected_ < 0.5 (or RTC > 0.9) are plotted in colors as correspondingly noted at the top of the graphic; listed on top are Sherlock p-values, with Sherlock p_corrected_ ≤ 0.05 highlighted in bold. For each additional gene in the region with an eQTL, the single eSNP with minimal eQTL p-value (“max-eQTL”) is marked by a black point (corresponding genes names are located above the chromosome marker bar). Locations of regional protein-coding genes and non-coding RNAs with gene symbols that did not bear any eQTL (either expressed genes without detected eQTL, or genes with below-threshold expression) are depicted in gray as denoted. Vertical dotted lines mark recombination hotspot boundaries, and horizontal dotted lines denote the thresholds for eQTL and GWAS significance, as well as the ceiling imposed for visualization purposes. Association betas (effect sizes) are plotted for SNP alleles associated with SCZ risk, with the gray band corresponding to increased risk of SCZ in the locus. The red bands mark the estimated direction and magnitude of the effect of the risk genotypes on expression of the corresponding gene (*FURIN* and *CLCN3*, respectively), where values above the bolded 0 line mark up-regulation (*CLCN3*) and below the line down-regulation (*FURIN*). **(B)** For each of *FURIN* and *CLCN3*, the underlying association of expression with SCZ risk allele (cis-eQTL) is plotted for the GWAS index SNP in the respective locus from (A), where the shape corresponds to diagnosis. Association betas and p-values, for eQTL and GWAS, are as listed.

CLCN3, SNAP91, and TSNARE1 are direct synaptic components, and CNTN4 and FURIN play roles in neurodevelopment. Specifically, CLCN3 (or ClC-3) is a brain-expressed chloride channel, where it appears to control fast excitatory glutamatergic transmission (*39*). SNAP91 is enriched in the presynaptic terminal of neurons where it regulates clathrin-coated vesicles, the major means of vesicle recycling at the presynaptic membrane. TSNARE1 plays key roles in docking, priming, and fusion of synaptic vesicles with the presynaptic membrane in neurons, thus synchronizing neurotransmitter release into the synaptic cleft. CNTN4 is a member of the contactin extracellular cell matrix protein family responsible for development of neurons including network plasticity (*40*). It plays a key role in olfactory axon guidance (*41*), and there is evidence for association of copy number variants overlapping *CNTN4* with autism (*42*). FURIN processes precursor proteins to mature forms, including brain-derived neurotrophic factor (BDNF) (*43*, *44*), a key molecule in brain development whose down-modulation has been hypothesized as related to schizophrenia (*45*), and *BDNF* and *FURIN* are up-regulated in astrocytes in response to stress (*43*).

The major histocompatibility complex (MHC / human leukocyte antigen / HLA) region is consistently most highly associated with SCZ, but it is a difficult region to dissect for causal variation because of its unusually high linkage disequilibrium and gene density (>200 DLPFC-expressed genes in chr6:25-36 Mb). Nevertheless, only five genes in this locus were ranked highly by Sherlock and passed evaluation for concordance of associations (data file 2): *C4A*, *HCG17*, *VARS2*, *HLA-DMB*, and *BRD2*. Consistent with recent work identifying structural variation of the *C4* genes as partly mediating the genetic MHC association, resulting in higher expression and perhaps driving pathological synapse loss in schizophrenia (*46*), we found a strong correlation between the risk alleles for SCZ and up-regulation of expression of *C4A* (complement component 4A; Spearman’s **ρ** = 0.66, p < 10^−16^).

## Functional dissection of genes highlighted by eQTL in common risk loci

Our results point to a number of genes worthy of follow-up, and we sought an assay that was rapid and amenable to over-and under-expression. Manipulation of zebrafish embryos fits these requirements, especially for evaluation of anatomical phenotypes of early development, such as head and brain size (or area). Perturbing expression of one or more genes in zebrafish has been used to identify genes contributing to neuropsychiatric disorders (*47*-*49*). Therefore, we asked whether suppression or overexpression of the corresponding gene within each of the five SCZ risk loci could identify key proteins that regulate brain development. To evaluate the four genes up-regulated by risk alleles in the GWAS loci, we injected 200pg of human capped mRNA encoding *TSNARE1*, *CNTN4*, *SNAP91*, or *CLCN3* in 1-8 cell stage embryos (n = 60 per experiment, at least two biological replicates performed). At 3 days post-fertilization (dpf), we assessed the area of the head that contains the forebrain and midbrain structures (Fig. 3A, B). Relative to control embryos, overexpression of *TSNARE1* or *CNTN4* resulted in a significant decrease in head size, 9.5% (p < 0.001) and 3.5% (p = 0.018), respectively, while *SNAP91* or *CLCN3* showed no statistically significant effect (Fig. 3A, B). Body length and somitic structures were similar across all embryos, suggesting that our observations were unlikely due to gross developmental delay. For *FURIN*, we sought to mimic the transcriptional down-regulation in human brains associated with SCZ risk. A reciprocal BLAST search of the zebrafish genome revealed a *FURIN* ortholog with two potential paralogs; both copies were expressed at ∼40-60 counts per million reads in mRNA from heads of 3 dpf zebrafish embryos (*50*). We depleted *furin_a*, the isoform most closely resembling the human ortholog, using a splice blocking morpholino (sbMO) that almost completely extinguished expression of the endogenous message by triggering the inclusion of intron 7 (Supplementary Fig. 7). Suppression of *furin_a* led to a 24% decrease in head size (Fig. 3A, B); this observation was replicated with a second sbMO targeting exon 5 (data not shown). Importantly, expression of human *FURIN* mRNA could rescue the phenotype induced by either morpholino, providing evidence for specificity (Supplementary Fig. 7E).

**Figure 3.**
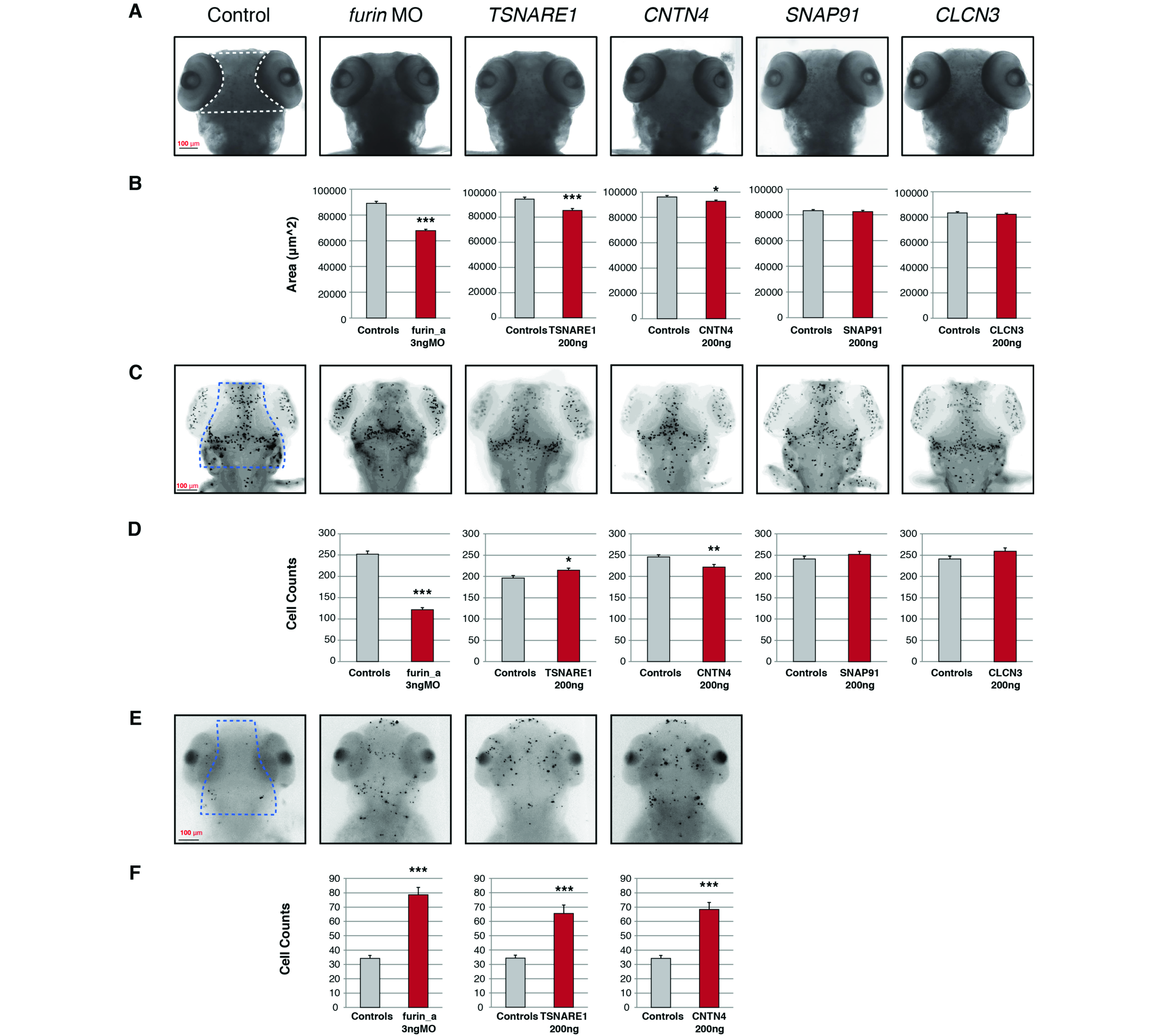
Neuroanatomical phenotypes upon suppression or overexpression of genes at SCZ risk loci. **(A)** Suppression of *furin_a* or overexpression of *TSNARE1* or *CNTN4* resulted in a smaller head size phenotype. Representative head size images per treatment condition are shown, and the area of the head quantified is depicted by the dashed white lines in the control image. **(B)** Quantification of head size phenotype in each treatment condition as compared to control embryos. **(C)** Representative images of PH3 staining assessing proliferation phenotypes. Dashed blue lines depict the area included in the quantification of cell counts. **(D)** Quantification of PH3-labeled cells with respect to each treatment condition. **(E)** Representative images of TUNEL staining per condition marking cells undergoing apoptosis. Area quantified is depicted within the dashed blue lines. **(F)** Cell counts of apoptotic cells in each treatment condition as compared to controls. Error bars are s.e., *p <0.05, **p<0.005, ***p<0.0005; MO - morpholino.

Given a potential role for *FURIN*, *TSNARE1*, and *CNTN4* during neurogenesis, we asked whether the decrease in head size could be attributed to changes in cell proliferation and/or apoptosis. Overexpression of *CNTN4* and suppression of *furin_a* led to a 9.8% (p = 0.003) and a 29.8% (p < 0.001) decrease, respectively, in proliferating cells marked by phospho-histone3 (PH3), and overexpression of *TSNARE1* led to a 9.5% increase (p = 0.018) in proliferating cells (n=20 per experiment; Fig. 3C, D). Next, we wondered how more proliferating cells nevertheless resulted in a smaller head size phenotype for the case of *TSNARE1*. To test the possibility that cells exiting cell cycle experience a higher apoptotic index, we performed TUNEL staining on injected embryos, and determined that modulation of all three target genes led to a significant increase in apoptotic cells in the head region corresponding to our head size measurements (n=20 per experiment; p < 0.001; Fig. 3E, F). Taken together, the data support the hypothesis that changes in *FURIN*, *TSNARE1*, and *CNTN4* expression levels induce subtle neuroanatomical variation in multiple brain regions.

Depletion of *furin* in our *in vivo* zebrafish model had the largest impact on head size. Thus we further tested the impact of *FURIN* knockdown in human neural progenitor cells (NPCs) capable of differentiating into mixed populations of post-mitotic neurons and astrocytes (*51*, *52*). Neurosphere outgrowth is a well-established neural migration assay measuring the distance NPCs migrate away from the neurosphere (*53*). NPCs were differentiated from human induced pluripotent stem cells (hiPSCs) reprogrammed from human fibroblasts using sendai viral vectors (*54*). Pairwise isogenic comparisons were conducted in 307 neurospheres from three independent unaffected controls. We measured migration of DAPI-positive nuclei from pLKO.1 non-hairpin-PURO control neurospheres (n = 147) and LV-*FURIN* shRNA-PURO (shRNA-*FURIN*) knockdown neurospheres (n = 160). *FURIN* knockdown in the hiPSC NPCs resulted in significantly decreased total radial migration for all three individuals (C1: 1.16-fold decrease, p < 0.0017; C2: 1.23-fold, p < 3 x 10^−6^; C3: 1.22-fold, p < 2 x 10^−6^) (Fig. 4).

**Figure 4.**
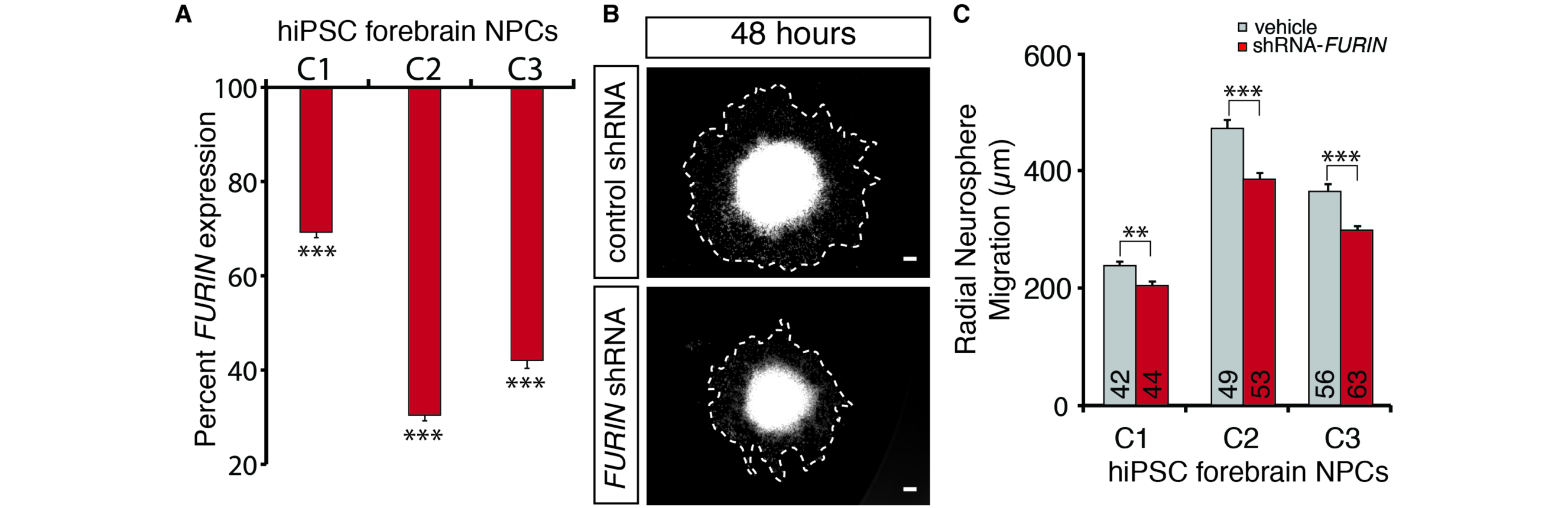
Decreasing *FURIN* expression in human NPCs perturbs neural migration. **(A)** *FURIN* expression reduction achieved by lentiviral (LV)-*FURIN* shRNA-PURO, relative to LV-non-hairpin-PURO control. **(B)** Representative images of the hiPSC NPC neurosphere outgrowth assay after 48 hours of migration, following transduction with LV-*FURIN* shRNA-PURO and LV-non-hairpin-PURO control. The average distance between the radius of the inner neurosphere (dense aggregate of nuclei) and outer circumference of cells (white dashed line) was calculated. DAPI-stained nuclei (blue), scale bar 100 μm. **(C)** Across hiPSC NPCs generated from three controls, average radial neurosphere migration following transduction with LV-*FURIN* shRNA-PURO (red bars) or LV-non-hairpin-PURO (gray bars). Error bars are s.e., *p<0.05, **p < 0.01, ***p < 0.001.

## Gene expression is subtly disrupted in schizophrenia

We next evaluated whether SCZ cases versus controls differed in their expression levels per gene. Following normalization of read counts for each gene, a weighted linear regression adjusting for known covariates was performed (Supplementary Figs. 2 and 4). Analysis of the distribution of p-values for the 16,423 genes was tested for a mixture of disease-associated and null distributions (*26*) and suggests that approximately 44% of genes are perturbed in SCZ; this excess of low p-values disappears when case and control labels are permuted (*55*). While polygenic inheritance, where many genes are affected but to a small degree (*7*) (*56*), could explain this result, treatment and environmental factors also likely play a role. Without imposing a threshold on the magnitude of fold change in mean expression between SCZ and controls, we find 693 genes to be differentially expressed after correction for multiple testing (FDR ≤ 5%), 332 up-regulated and 361 down-regulated (Fig. 5A, data file 3). All had modest fold changes (Fig. 5B), with a mean of 1.09 and range 1.03-1.33 (inverting down-regulated expression ratios). As expected, hierarchical clustering of the differentially expressed genes showed case-control distinctions but were independent of institution, sex, age at death, ethnicity, and RIN (Fig. 5A). We examined differential expression in an independent sample, the NIMH Human Brain Collection Core (HBCC), which generated DLPFC gene expression data using Illumina HumanHT-12_V4 Beadchip microarrays from 131 SCZ cases and 176 controls. Though these arrays differ from RNA-seq in their capture features, there was high correlation of test statistics for differential expression in CMC compared to HBCC for the differentially expressed genes also present in the HBCC data (480 of 693), Pearson correlation r = 0.58 (p < 10^−16^); the correlation remains high (r = 0.28, p < 10^−16^) across all 10,928 genes common to both platforms after QC (Fig. 5C).

**Figure 5.**
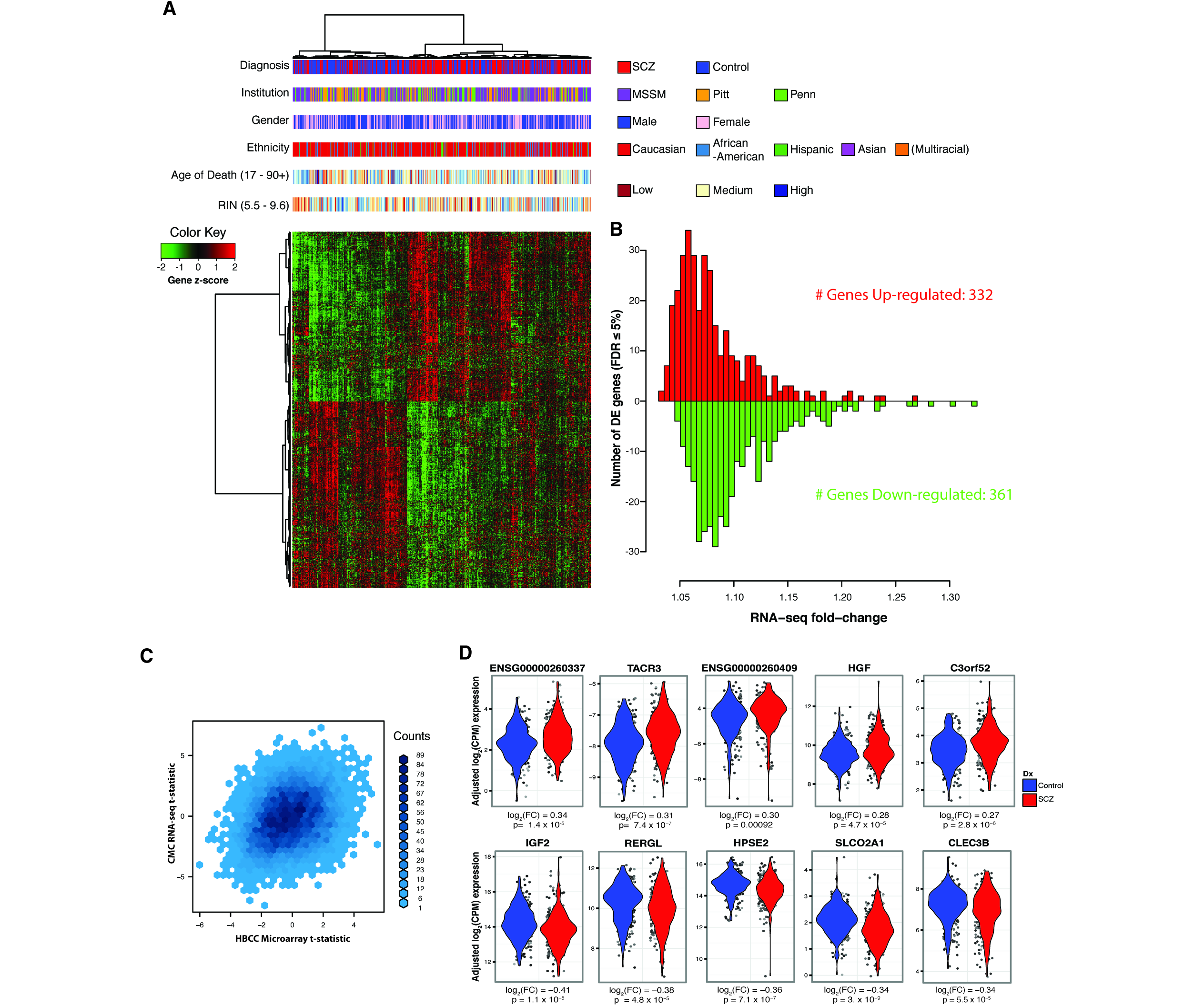
Differential expression between schizophrenia cases and controls in the DLPFC. **(A)** For the N = 693 genes differentially expressed at FDR ≤ 5%, bivariate clustering of individuals (columns) and genes (rows) depicts the case-control differences, as marked by the red-blue horizontal colorbar at top (’Diagnosis’). The expression for an individual (converted to a z-score per gene) is marked in red if it is higher than other individuals, and green if lower than others; thus, the top half of the plot consists of genes up-regulated in cases versus controls (green in top left; red in top middle), and the bottom half of down-regulated genes (red in bottom left; green in bottom middle). In addition to the horizontal colorbar marking case-control status for each sample, additional colorbars denote brain bank (’Institution’), gender, reported ancestry (’Ethnicity’), age of death, and RNA quality (’RIN’); note that the latter two use a continuous-values color scale (with low, medium, and high as colored), and minimum and maximum values are given in parentheses. **(B)** Distribution of fold-change of differential expression for 693 differentially expressed genes. Case:control fold-changes for up-regulated genes are plotted in red pointing upwards, and control:case fold-changes for down-regulated genes in green facing down. **(C)** Binned density scatter plot comparing the t-statistics for case versus control differential expression between the independent HBCC replication cohort assayed on microarrays and the CommonMind RNA-seq data; correlation between the statistics is 0.28. **(D)** For the 10 significantly differentially expressed genes with the largest fold changes (5 up-and 5 down-regulated), the distributions of normalized and adjusted gene expression in cases (red) versus controls (blue).

The differential expression observed here is smaller than that reported in earlier studies (data file 1), but it is consistent with plausible models for average differential gene expression and the polygenic inheritance of SCZ (Supplementary Text, with meta-analysis of earlier studies Supplementary Fig. 8). Consider, for example, a gene for which the major determinant of differential expression is the case-control difference in allele frequency at an eQTL SNP. For that gene, the expected magnitude of differential expression fold change will be on the order of the allele frequency differences seen in the recent large Psychiatric Genomic Consortium SCZ genetic association study (∼1-2%) (*7*), precisely what is observed in the CMC data. Such modeling can also explain the difference between earlier studies and CMC results; because earlier studies tend to be far smaller in sample size, their larger differential expression is consistent with either the well-known “Winner’s Curse” (*57*) or false positives that may occur in smaller samples. Finally, our results imply a need for thousands of samples to ensure 80% statistical power to observe differential expression between cases and controls for the genes implicated at SCZ-associated eQTL, e.g., the five genes of interest above.

The most highly up-regulated protein-coding gene is tachykinin receptor 3 (*TACR3*, NK_3_ receptor, 1.24-fold, Fig. 5D). NK3 antagonists have been tested in SCZ and other CNS diseases (*58*). Moreover, rat and human studies have suggested a role for the NK_3_ receptor in memory and cognition (*59*), both key impairments of schizophrenia (*60*). Insulin-like growth factor 2 (*IGF2*), the most strongly down-regulated gene (1.33-fold, Fig. 5D), can rescue neurogenesis and cognitive deficits in certain mouse models of schizophrenia (*61*). Also included among the top 100 differentially expressed genes are the alpha 5 subunit of the GABA A receptor (*GABRA5*) (*62*) and calbindin (*CALB1*) (*63*), genes previously reported as differentially expressed in cortical tissue from schizophrenia patients, suggesting GABAergic interneuron dysfunction (*64*).

We identified 239 isoforms differentially expressed between SCZ cases and controls: 94 up-regulated and 145 down-regulated. These isoforms derive from 223 genes, which are enriched, as expected, for overlap with the 693 differentially expressed genes (p = 2 x 10^−131^, Fisher’s exact test), and 136 are differentially expressed at both the gene and isoform levels (Supplementary Fig. 9). No obvious unifying biological theme emerges from this set of genes and isoforms on the basis of pathway enrichment analysis (data file 4). An assessment of the impact of age at death or cell type proportions suggests that these variables do not explain significant differential expression (Supplementary Fig. 10). Although analyses of experiments performed using either monkeys or rodents indicate that genes whose expression are affected by antipsychotics are often the same as those we find altered in individuals with SCZ, the impact of antipsychotic drugs nevertheless tends to be significantly in the opposite direction of that observed in the SCZ subjects (Supplementary Table 2). Thus, our analyses find that genes highlighted by the contrast of SCZ cases versus control subjects do not largely trace their differential expression to antipsychotic medications, although intriguingly they do suggest a mechanism for the efficacy of these drugs (*65*).

## Brain co-expression networks capture SCZ associations

Coordinated expression of genes is critical to brain development and function. One expectation of polygenic inheritance of disease is that this coordination may be subtly altered in individuals with SCZ. To assess this, we applied weighted gene co-expression network analysis (WGCNA) (*66*) to the matrix of pairwise gene co-expression values. WGCNA recovers a network that consists of nodes (genes) and edges connecting nodes (i.e., the degree of co-expression for a pair of genes, measured as their correlation after transformation by raising the value to a power β that results in an overall scale-free topology). WGCNA divides the network into subnetworks called modules, or clusters of genes with more highly correlated expression.

We constructed gene co-expression networks separately from control individuals and SCZ cases (data file 5), since we wished to assess disease-dependent changes in co-expression for modules of interest (*27*). The co-expression network generated from the controls consisted of 35 modules each containing between 30 and 1,900 genes, along with ∼3,600 unclustered genes (data file S5). Four modules stand out in harboring an excess of differentially expressed genes (Fig. 6A, data file 6). Of these, however, only one (M2c) shows association with differential expression (OR = 2.3, p = 1 x 10^−13^) and multiple prior genetic associations with SCZ; the latter encompasses genes in GWAS loci (FE [fold-enrichment] = 1.36, p = 0.04), rare CNV (FE = 1.52, p = 0.051), and rare nonsynonymous variants (FE = 1.18, p = 2 x 10^−4^) (Supplementary table 3). Given its apparent relevance to SCZ risk, we tested if the co-expression pattern for M2c was perturbed in SCZ samples relative to controls. We used two categories of network-based preservation statistics: (a) testing whether highly connected nodes in a module remain as highly connected (“density”), or (b) testing for differences in the overall connectivity pattern in a module (“connectivity”). The M2c module exhibits a loss of density in the SCZ cases (permutation *Z* = −1.79, one-tailed p = 0.037, Fig. 6B) but no loss of connectivity. The loss of density replicates in the HBCC cohort (*Z* = −3.02, p = 0.003), indicating that the regulatory coordination of genes in this module is disrupted in SCZ. The dysregulation of M2c in SCZ is not due to medication effect or clinical and technical confounds (See Supplement).

**Figure 6.**
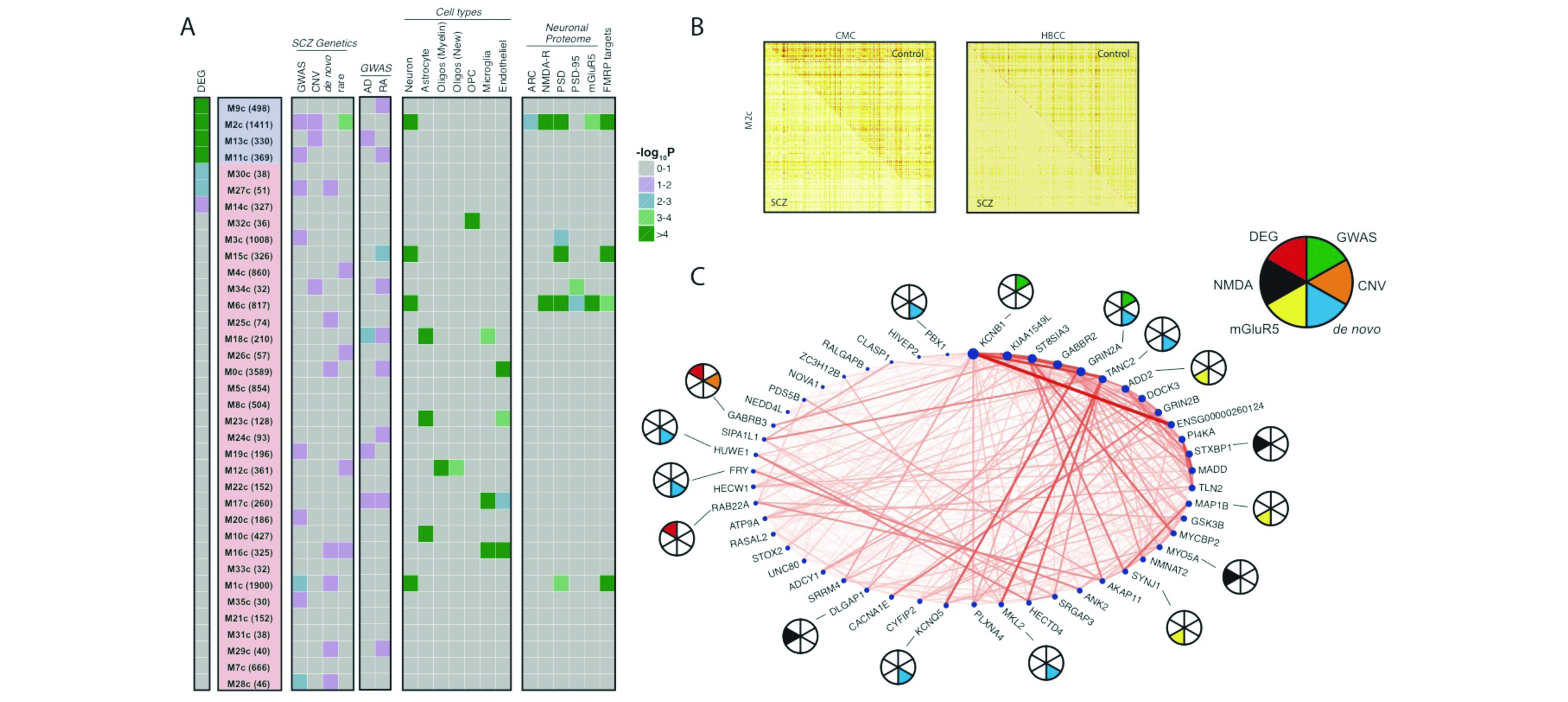
Co-expression network analysis in control DLPFC samples. **(A)** Control-derived modules were ranked by enrichment with differentially expressed genes (DEG); number of genes in each module is given in parentheses. Among the 4 modules with strongest overlap (marked in blue), only the M2c module genes are strongly enriched for multiples lines of prior genetic evidence. The enrichment of each module with SCZ genetics, cell type-specific markers, neuronal proteome sets, and FMRP targets is depicted at right. Note the lack of enrichment of M2c with common variants for Alzheimer’s disease (AD) and rheumatoid arthritis (RA) **(B)** Topological overlap matrix of the differentially connected M2c module in controls (upper right triangle) and SCZ cases (lower left triangle) in the CMC (left) and HBCC (right) cohorts. **(C)** Circle plot showing connection strengths for the top 50 hub genes of the M2c module, where node size corresponds to intramodular connectivity and nodes are ordered clockwise based on connectivity. Pie chart: SCZ susceptibility genes based on GWAS PGC2-SCZ (green), CNV (orange) or *de novo* (cyan) studies; Genes that belong in the NMDA (black) or mGluR5 (yellow) signalling pathway; Genes that are differentially expressed in schizophrenia vs. controls at FDR ≤ 5% (red).

Consistent with prior studies of the brain transcriptome (*27*, *67*-*70*), we find gene co-expression to be organized into modules of distinct cellular and functional categories (data file 7). In particular, the M2c module is enriched for multiple categories, including axon guidance, postsynaptic membrane, transmission across chemical synapses, and voltage-gated potassium channel complexes (Fig. 6C). Gene sets identified in prior genetic studies that highlighted certain neurobiological functions are also enriched in the M2c module, including the activity-regulated cytoskeleton-associated (ARC) protein complex, targets of fragile X mental retardation protein (FMRP), neuronal markers, post-synaptic density (PSD) proteins, and NMDA receptors (Fig. 6A). Overall, our results point to the M2c module of ∼1400 genes that possess functions related to synaptic transmission as being enriched for differential expression, overlapping SCZ genetic signal, and with some genes having less dense co-expression in SCZ cases.

## Conclusions

The findings reported here by the CommonMind Consortium (CMC) represent a unique resource to understand brain function, basic neuroscience, and brain diseases at the molecular level. They include a comprehensive compilation of gene expression patterns, together with intensive evaluation of eQTLs across the genome. The expertise and support to produce and analyze these data required a consortium of brain banks, pharmaceutical companies, a foundation, academic centers, and the NIMH, and this work represents the first phase of our ongoing project. All results are available through the CommonMind Knowledge Portal (www.synapse.org/CMC) with a searchable database of eQTLs and other visualizations (shiny.synapse.org/users/ssiebert/cmc_eqtl_query/). Both alone, and in combination with other datasets such as GTEx, the CMC data will empower future studies of disease and the brain.

We used these data to understand more about the genetics and molecular etiology of SCZ. Our analyses had two fundamental goals: to identify mechanisms that underlie genetic risk, and to describe differences in gene expression and co-expression related to disease. By intersecting transcriptomics and genetics, we elucidated important aspects of the genetic control of transcription and found that 20 of the 108 SCZ GWAS risk variants alter expression of one or more genes. Prior analyses using older brain eQTL datasets pointed to only three such associations (*3*). We demonstrated that experimental manipulation of three of five genes for which GWAS variants alter expression had an impact on neuroanatomical and developmental attributes in model systems. We also detected replicable differences in gene expression in SCZ that point to subtle but broad disruption in transcription, which is consistent with the polygenic nature of SCZ genetics and possibly other disease-related factors. This study paves the way for connecting genetic influences on cellular function with changes in macroscopic circuits of the brain that may ultimately lead to disease.

## ONLINE METHODS

Postmortem human brain samples were collected for schizophrenia or schizoaffective disorder (n=258) cases, control subjects (n=279), and cases with affective disorders (n=55), from three brain banks: Mount Sinai NIH Brain Bank and Tissue Repository, University of Pittsburgh NIH NeuroBioBank Brain and Tissue Repository, and University of Pennsylvania Brain Bank of Psychiatric illnesses and Alzheimer’s Disease Core Center. DNA and RNA were extracted from dorsolateral prefontal cortex (Brodmann areas 9/46) tissue. DNA was genotyped using the Illumina Infinium HumanOmniExpressExome chip, and QC was performed using PLINK. rRNA was depleted from total RNA using Ribo-Zero Magnetic Gold kit, a DNA sequencing library was prepared using the TruSeq RNA Sample Preparation Kit, and the library was subjected to paired-end sequencing on Illumina HiSeq sequencers. The RAPiD pipeline used TopHat for alignment of reads to human reference genome hg19 guided by Ensembl v70 gene models, followed by gene-level quantification using HTSeq and isoform-level abundance estimation using MISO. The gene-level expression matrix was normalized to log(counts per million) using voom, known covariates (encompassing sample ascertainment and quality, experimental parameters, and individual ancestry) were selected for adjustment, surrogate variables were extracted to explain additional variance (for eQTL only), and covariates were adjusted using linear modeling, with voom-derived regression weights. Expression quantitative trait loci (eQTL) were detected across all genetically-inferred Caucasian samples using MatrixEQTL, controlling for sample ancestry and diagnosis. eQTL associations were compared using Sherlock to genome-wide associations (GWAS) for schizophrenia, to functionally fine-map associated disease loci for causal genes. Sherlock disease gene predictions were additionally filtered for strict concordance between the genetic association with expression and disease. Overexpression or morpholino-driven suppression of expression in zebrafish was performed for 5 genes prioritized by Sherlock, followed by assessment of differences in head size, neural proliferation, and apoptosis. To test the effect of *FURIN* knockdown, neural migration was assayed in neural progenitor cells (NPCs) differentiated from human induced pluripotent stem cells (hiPSCs) reprogrammed from human fibroblasts from control individuals. Limma-based linear regression was used for schizophrenia case-control differential expression analysis, and differentially expressed genes were tested for enrichment in schizophrenia genetics, and with Bonferroni multiple test correction for other gene sets using standard approaches. Gene co-expression networks were constructed using WCGNA separately for schizophrenia cases and controls, co-expression modules were extracted, and case and control modules were contrasted for differential co-expression using a permutation-based approach. Module genes were tested for enrichment in disease genetics and other gene sets.

## AUTHOR CONTRIBUTIONS

PR, JSJ, KT, AC, REG, CH, DAL, VH, BKL and JDB contributed to sample collection. SEH contributed monkey brain tissue.

MF, PR, SKS, DHK, TMP, DMR, KKD, PFS, AC, MAP, JDB, ED, BD and PS contributed to the writing of this manuscript.

MF, PR, SKS, DMR, HRS, KKD, JMD, AC, SMP, LAS, LMM, HT, DAL, MAP, JDB, EES, KH,

KJB, NK, BD and PS contributed to experimental and study design and planning analytical strategies.

AC, LAS, HT, DAL, BKL, JDB, EES, KH, ED, BD and PS contributed the funding of this work.

MF, PR, SKS, JSJ, DHK, TMP, DMR, HRS, LLK, RK, DP, ZHG, AC, KKD, AB, CL, BR, EAS,

TH, JFF, YW, JD, BAL, TR, JZ, BZ, PFS, SMP, EES, ED, BD and PS contributed to data analyses.

ECO, AT, MP, KJB and NK contributed to the model system experiments.

TR, DAB, PLD contributed the ROS/MAP data.

MCM, JMD, AC, LAS, LMM, HT, REG, CH, DAL, MAP, BKL, JDB, EES, KH, ED, BD and PS contributed to the management and leadership of the CommonMind Consortium.

## ACKNOWLEDGMENTS

We thank the patients and families who donated material for these studies. We thank Thomas Lehner for organizational and intellectual support. We thank Xin He for helpful discussions regarding Sherlock, Joe Scarpa for help running and interpreting WGCNA, Laurent Essioux for support in establishing and managing interactions with the Consortium, and Alessandro Bertolino and Anirvan Ghosh for continuous encouragement. Data were generated as part of the CommonMind Consortium supported by funding from Takeda Pharmaceuticals Company Limited, F. Hoffman-La Roche Ltd and grants R01MH093725-02S1 (JB), P50MH066392 (JB), R01MH097276 (PS, ES), R01MH075916 (CGH), P50MH096891 (CGH), P50MH084053-S1 (DAL), R37MH057881 (BD) and R37MH057881S1 (BD), R01MH085542-S1 (PS), U01MH096296-S2 (PS), HHSN271201300031C (VH), VA VISN3 MIRECC (VH), P50MH066392 (JDB), NIMH Intramural program (BKL), R01MH101454 (KJB), New York Stem Cell Foundation (KJB), the Silvio O Conte Center grant P50MH094268 (NK), NARSAD (ECO), NARSAD Young Investigator (EAS), and the Stanley Medical Research Foundation and NIMH-R01MH074313 (SEH). Brain tissue for the study was obtained from the following brain bank collections: the Mount Sinai NIH Brain and Tissue Repository, the University of Pennsylvania Alzheimer’s Disease Core Center, the University of Pittsburgh Brain Tissue Donation Program, the NIMH Human Brain Collection Core, and Wake Forest University. CMC Leadership: Pamela Sklar, Joseph Buxbaum (Icahn School of Medicine at Mount Sinai), Bernie Devlin, David Lewis (University of Pittsburgh), Raquel Gur, Chang-Gyu Hahn (University of Pennsylvania), Keisuke Hirai, Hiroyoshi Toyoshiba (Takeda Pharmaceuticals Company Limited), Enrico Domenici, Laurent Essioux (F. Hoffman-La Roche Ltd), Lara Mangravite, Mette Peters (Sage Bionetworks), Thomas Lehner, Barbara Lipska (NIMH). The GTEx data used for the analyses described herein were obtained from the GTEx Portal (www.gtexportal.org), corresponding to dbGaP accession number phs000424.v6.p1.

## REFERENCES

1 J. McGrath, S. Saha, D. Chant, J. Welham, Schizophrenia: a concise overview of incidence, prevalence, and mortality. Epidemiologic reviews 30, 67–76 (2008).

2 G. Kirov, CNVs in neuropsychiatric disorders. Hum Mol Genet 24, R45–49 (2015).

3 C. Schizophrenia Working Group of the Psychiatric Genomics, Biological insights from 108 schizophrenia-associated genetic loci. Nature 511, 421–427 (2014).

4 S. M. Purcell et al., Common polygenic variation contributes to risk of schizophrenia and bipolar disorder. Nature 460, 748–752 (2009).

5 S. M. Purcell et al., A polygenic burden of rare disruptive mutations in schizophrenia. Nature 506, 185–190 (2014).

6 R. Fisher, The correlation between relatives on the supposition of Mendelian inheritance. Trans Roy Soc Edinb 52, 399–433 (1918).

7 S. W. G. o. t. P. G. Consortium, Biological insights from 108 schizophrenia-associated genetic loci. Nature 511, 421–427 (2014).

8 T. Walsh et al., Rare structural variants disrupt multiple genes in neurodevelopmental pathways in schizophrenia. Science 320, 539–543 (2008).

9 M. Fromer et al., De novo mutations in schizophrenia implicate synaptic networks. Nature 506, 179–184 (2014).

10 S. Horvath, Z. Janka, K. Mirnics, Analyzing schizophrenia by DNA microarrays. Biol Psychiatry 69, 157–162 (2011).

11 M. Mistry, J. Gillis, P. Pavlidis, Meta-analysis of gene coexpression networks in the post-mortem prefrontal cortex of patients with schizophrenia and unaffected controls. BMC neuroscience 14, 105 (2013).

12 Z. Wang, M. Gerstein, M. Snyder, RNA-Seq: a revolutionary tool for transcriptomics. Nature reviews. Genetics 10, 57–63 (2009).

13 R. Hitzemann et al., Introduction to sequencing the brain transcriptome. International review of neurobiology 116, 1–19 (2014).

14 J. Mudge et al., Genomic convergence analysis of schizophrenia: mRNA sequencing reveals altered synaptic vesicular transport in post-mortem cerebellum. PLoS One 3, e3625 (2008).

15 J. Q. Wu et al., Transcriptome sequencing revealed significant alteration of cortical promoter usage and splicing in schizophrenia. PLoS One 7, e36351 (2012).

16 K. C. Huang, K.C. Yang, H. Lin, T.T. Tsao, S. A. Lee, Transcriptome alterations of mitochondrial and coagulation function in schizophrenia by cortical sequencing analysis. BMC genomics 15 Suppl 9, S6 (2014).

17 R. Kohen, A. Dobra, J. H. Tracy, E. Haugen, Transcriptome profiling of human hippocampus dentate gyrus granule cells in mental illness. Translational psychiatry 4, e366 (2014).

18 Y. Xiao et al., The DNA methylome and transcriptome of different brain regions in schizophrenia and bipolar disorder. PLoS One 9, e95875 (2014).

19 J. A. Miller et al., Transcriptional landscape of the prenatal human brain. Nature 508, 199–206 (2014).

20 A. E. Jaffe et al., Developmental regulation of human cortex transcription and its clinical relevance at single base resolution. Nature neuroscience 18, 154–161 (2015).

21 G. R. Abecasis et al., An integrated map of genetic variation from 1,092 human genomes. Nature 491, 56–65 (2012).

22 A. A. Shabalin, Matrix eQTL: ultra fast eQTL analysis via large matrix operations. Bioinformatics 28, 1353–1358 (2012).

23 J. B. Veyrieras et al., High-resolution mapping of expression-QTLs yields insight into human gene regulation. PLoS Genet 4, e1000214 (2008).

24 Human genomics. The Genotype-Tissue Expression (GTEx) pilot analysis: multitissue gene regulation in humans. Science 348, 648–660 (2015).

25 I. Dunham et al., An integrated encyclopedia of DNA elements in the human genome. Nature 489, 57–74 (2012).

26 J. D. Storey, R. Tibshirani, Statistical methods for identifying differentially expressed genes in DNA microarrays. Methods Mol Biol 224, 149–157 (2003).

27 B. Zhang et al., Integrated systems approach identifies genetic nodes and networks in late-onset Alzheimer’s disease. Cell 153, 707–720 (2013).

28 C. Colantuoni et al., Temporal dynamics and genetic control of transcription in the human prefrontal cortex. Nature 478, 519–523 (2011).

29 J. R. Gibbs et al., Abundant quantitative trait loci exist for DNA methylation and gene expression in human brain. PLoS Genet 6, e1000952 (2010).

30 A. Ramasamy et al., Genetic variability in the regulation of gene expression in ten regions of the human brain. Nature neuroscience 17, 1418–1428 (2014).

31 Y. Kim et al., A meta-analysis of gene expression quantitative trait loci in brain. Translational psychiatry 4, e459 (2014).

32 F. A. Wright et al., Heritability and genomics of gene expression in peripheral blood. Nat Genet 46, 430–437 (2014).

33 P. Roussos et al., A role for noncoding variation in schizophrenia. Cell Rep 9, 1417–1429 (2014).

34 A. L. Richards et al., Schizophrenia susceptibility alleles are enriched for alleles that affect gene expression in adult human brain. Mol Psychiatry 17, 193–201 (2012).

35 G. Trynka et al., Chromatin marks identify critical cell types for fine mapping complex trait variants. Nat Genet 45, 124–130 (2013).

36 G. Trynka et al., Disentangling the Effects of Colocalizing Genomic Annotations to Functionally Prioritize Non-coding Variants within Complex-Trait Loci. Am J Hum Genet 97, 139–152 (2015).

37 X. He et al., Sherlock: detecting gene-disease associations by matching patterns of expression QTL and GWAS. Am J Hum Genet 92, 667–680 (2013).

38 P. L. De Jager et al., Alzheimer’s disease: early alterations in brain DNA methylation at ANK1, BIN1, RHBDF2 and other loci. Nature neuroscience 17, 1156–1163 (2014).

39 R. E. Guzman, A. K. Alekov, M. Filippov, J. Hegermann, C. Fahlke, Involvement of ClC-3 chloride/proton exchangers in controlling glutamatergic synaptic strength in cultured hippocampal neurons. Frontiers in cellular neuroscience 8, 143 (2014).

40 Y. Shimoda, K. Watanabe, Contactins: emerging key roles in the development and function of the nervous system. Cell adhesion & migration 3, 64–70 (2009).

41 T. Kaneko-Goto, S. Yoshihara, H. Miyazaki, Y. Yoshihara, BIG-2 mediates olfactory axon convergence to target glomeruli. Neuron 57, 834–846 (2008).

42 J. T. Glessner et al., Autism genome-wide copy number variation reveals ubiquitin and neuronal genes. Nature 459, 569–573 (2009).

43 Y. Chen, J. Zhang, M. Deng, Furin mediates brain-derived neurotrophic factor upregulation in cultured rat astrocytes exposed to oxygen-glucose deprivation. Journal of neuroscience research 93, 189–194 (2015).

44 N. Adachi, T. Numakawa, M. Richards, S. Nakajima, H. Kunugi, New insight in expression, transport, and secretion of brain-derived neurotrophic factor: Implications in brain-related diseases. World journal of biological chemistry 5, 409–428 (2014).

45 Q. Yuan et al., Regulation of Brain-Derived Neurotrophic Factor Exocytosis and Gamma-Aminobutyric Acidergic Interneuron Synapse by the Schizophrenia Susceptibility Gene Dysbindin-1. Biol Psychiatry, (2015).

46 A. Sekar et al., Schizophrenia risk from complex variation of complement component 4. Nature 530, 177–183 (2016).

47 K. Mishra-Gorur et al., Mutations in KATNB1 cause complex cerebral malformations by disrupting asymmetrically dividing neural progenitors. Neuron 84, 1226–1239 (2014).

48 C. Golzio et al., KCTD13 is a major driver of mirrored neuroanatomical phenotypes of the 16p11.2 copy number variant. Nature 485, 363–367 (2012).

49 C. M. Carvalho et al., Dosage changes of a segment at 17p13.1 lead to intellectual disability and microcephaly as a result of complex genetic interaction of multiple genes. Am J Hum Genet 95, 565–578 (2014).

50 G. Borck et al., BRF1 mutations alter RNA polymerase III-dependent transcription and cause neurodevelopmental anomalies. Genome research 25, 155–166 (2015).

51 K. J. Brennand et al., Modelling schizophrenia using human induced pluripotent stem cells. Nature, (2011).

52 A. Topol, N. N. Tran, K. J. Brennand, A guide to generating and using hiPSC derived NPCs for the study of neurological diseases. Journal of visualized experiments: JoVE, e52495 (2015).

53 C. Delaloy et al., MicroRNA-9 coordinates proliferation and migration of human embryonic stem cell-derived neural progenitors. Cell Stem Cell 6, 323–335 (2010).

54 I. S. Lee et al., Characterization of molecular and cellular phenotypes associated with a heterozygous CNTNAP2 deletion using patient-derived hiPSC neural cells. NPJ Schizophrenia 1, 15019 (2015).

55 S. Mostafavi et al., Type I interferon signaling genes in recurrent major depression: increased expression detected by whole-blood RNA sequencing. Mol Psychiatry 19, 1267–1274 (2014).

56 Gottesman, II, J. Shields, A polygenic theory of schizophrenia. Proc Natl Acad Sci U S A 58, 199–205 (1967).

57 R. Xiao, M. Boehnke, Quantifying and correcting for the winner’s curse in genetic association studies. Genetic epidemiology 33, 453–462 (2009).

58 L. A. Dawson, R. A. Porter, Progress in the development of neurokinin 3 modulators for the treatment of schizophrenia: molecule development and clinical progress. Future medicinal chemistry 5, 1525–1546 (2013).

59 M. A. de Souza Silva et al., Neurokinin3 receptor as a target to predict and improve learning and memory in the aged organism. Proc Natl Acad Sci U S A 110, 15097–15102 (2013).

60 R. S. Keefe, P. D. Harvey, Cognitive impairment in schizophrenia. Handbook of experimental pharmacology, 11–37 (2012).

61 Y. Ouchi et al., Reduced adult hippocampal neurogenesis and working memory deficits in the Dgcr8-deficient mouse model of 22q11.2 deletion-associated schizophrenia can be rescued by IGF2. J Neurosci 33, 9408–9419 (2013).

62 F. Impagnatiello et al., A decrease of reelin expression as a putative vulnerability factor in schizophrenia. Proc Natl Acad Sci U S A 95, 15718–15723 (1998).

63 S. J. Fung, S. G. Fillman, M. J. Webster, C. Shannon Weickert, Schizophrenia and bipolar disorder show both common and distinct changes in cortical interneuron markers. Schizophr Res 155, 26–30 (2014).

64 T. Sakai et al., Changes in density of calcium-binding-protein-immunoreactive GABAergic neurons in prefrontal cortex in schizophrenia and bipolar disorder. Neuropathology: official journal of the Japanese Society of Neuropathology 28, 143–150 (2008).

65 L. Carboni, E. Domenici, Proteome effects of antipsychotic drugs: Learning from preclinical models. Proteomics. Clinical applications, (2015).

66 B. Zhang, S. Horvath, A general framework for weighted gene co-expression network analysis. Statistical applications in genetics and molecular biology 4, Article17 (2005).

67 I. Voineagu et al., Transcriptomic analysis of autistic brain reveals convergent molecular pathology. Nature 474, 380–384 (2011).

68 A. Torkamani, B. Dean, N. J. Schork, E. A. Thomas, Coexpression network analysis of neural tissue reveals perturbations in developmental processes in schizophrenia. Genome research 20, 403–412 (2010).

69 M. C. Oldham et al., Functional organization of the transcriptome in human brain. Nature neuroscience 11, 1271–1282 (2008).

70 P. Roussos, P. Katsel, K. L. Davis, L. J. Siever, V. Haroutunian, A system-level transcriptomic analysis of schizophrenia using postmortem brain tissue samples. Arch Gen Psychiatry 69, 1205–1213 (2012).

71 K. Wang, M. Li, H. Hakonarson, ANNOVAR: functional annotation of genetic variants from high-throughput sequencing data. Nucleic acids research 38, e164 (2010).

